# Gamma-Band Correlations in Primary Visual Cortex

**DOI:** 10.1101/339184

**Authors:** X. Liu, P. Sanz-Leon, P. A. Robinson

**Author notes:** Corresponding Author (X.Liu).

## Abstract

Neural field theory is used to quantitatively analyze the two-dimensional spatiotemporal correlation properties of gamma-band (30 - 70 Hz) oscillations evoked by stimuli arriving at the primary visual cortex (V1), and modulated by patchy connectivities that depend on orientation preference (OP). Correlation functions are derived analytically under different stimulus and measurement conditions. The predictions reproduce a range of published experimental results, including the existence of two-point oscillatory temporal cross-correlations with zero time-lag between neurons with similar OP, the influence of spatial separation of neurons on the strength of the correlations, and the effects of differing stimulus orientations.

**Graphical Abstract:** 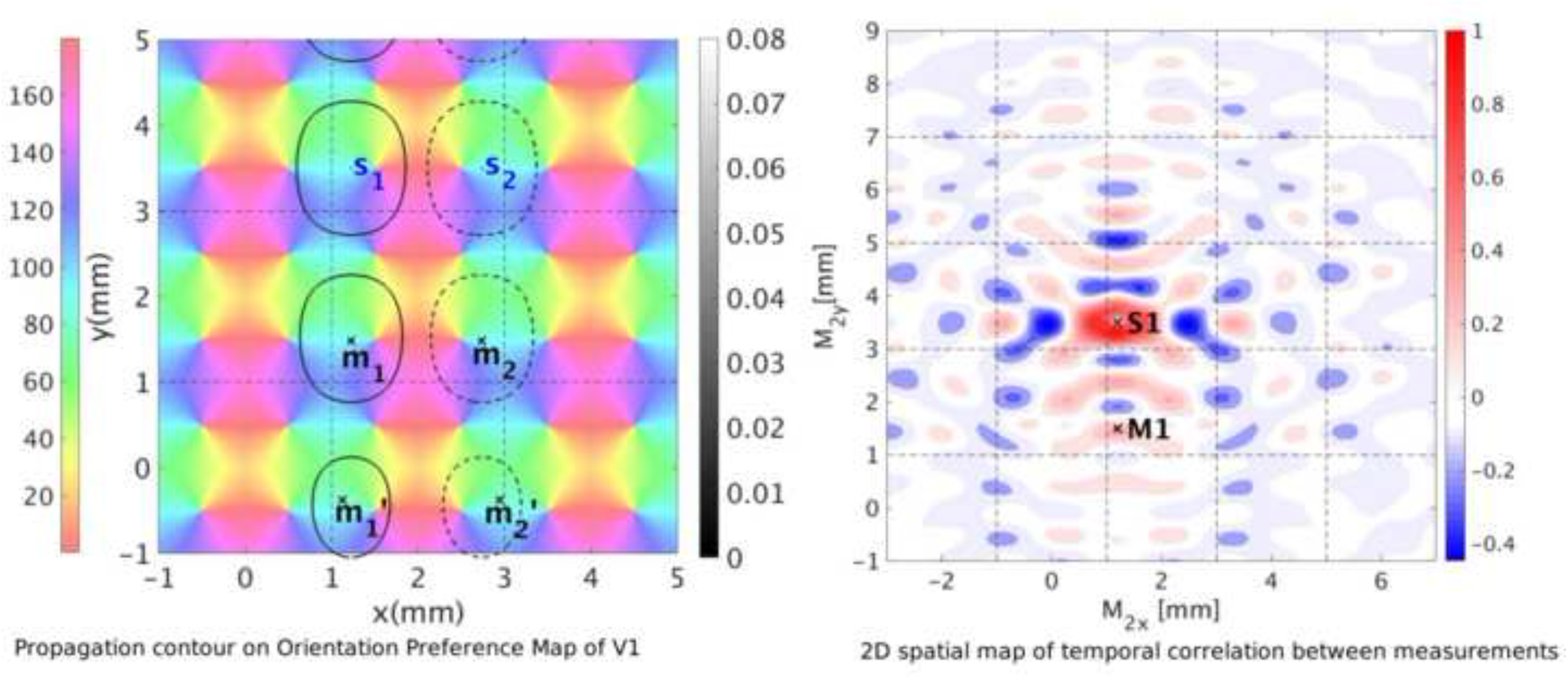

**Highlights:**

- Incorporate orientation preference map into patchy connectivities of neurons in V1.
- Generalize spatiotemporal correlation function of neural activity to 2D spatially.
- Reproduce experimental results: synchronization of neural activities in gamma band.
- Temporal correlation between 2 measurements decreases with increasing separation.
- Predict 2D spatial map of temporal correlation strength between 2 measurement sites.

## 1 Introduction

The primary visual cortex (V1) is the first cortical area to process visual inputs that arrive from the retina via the lateral geniculate nucleus of the thalamus (LGN) and passes the processed signals forward to higher visual areas, and back to the LGN. The feed-forward visual pathway from the eyes to V1 is such that the neighboring cells in V1 respond to neighboring regions of the retina (Schiller and Tehovnik, 2015). V1 can be approximated as a two-dimensional layered sheet (Tovée, 1996). Neurons that span vertically through multiple layers of V1 form a functional cortical column, and these neurons respond most strongly to a preferred stimulus orientation, right or left eye, direction of motion, and other feature preferences. Thus, various features of the visual inputs are mapped to V1 in different ways. These maps are overlaid such that a single neural cell responds to several features and all preferences within a given visual field are mapped to a small region of V1. Such an area of V1 is sometimes termed a hypercolumn, which corresponds to a particular visual field in the overall field of vision (Hubel and Wiesel, 1962, 1974; Miikkulainen et al., 2005).

A prominent feature of V1 is the presence of ocular dominance (OD) stripes, which reflect the fact that left- and right-eye inputs are mapped to alternating stripes ~ 1 mm wide, with each hypercolumn including left- and right-eye OD regions. Orientation preference (OP) of neurons for particular edge orientations in a visual field is mapped to each hypercolumn such that neurons with particular OP are located adjacent to one another and OP spans the range from 0° to 180°. Typically, OP varies with azimuth relative to a center, or singularity, in an arrangement called a pinwheel. The OP angle in each pinwheel rotates either clockwise (negative pinwheel) or counterclockwise (positive pinwheel), and neighboring pinwheels have opposite signs (Blasdel, 1992; Braitenberg and Braitenberg, 1979; Götz, 1987, 1988; Swindale, 1996). Hence, a hypercolumn must have left and right OD stripes with positive and negative pinwheels in each, as suggested by Bressloff and Cowan (2002) and Veltz et al. (2015). In Figs 1(a) and (b) we illustrate a negative pinwheel and a positive pinwheel, respectively, while Figs 1(c) and (d) show a hypercolumn and an array of hypercolumns, respectively; in the latter case the hypercolumn is the unit cell of the lattice and the schematic resembles maps reconstructed from *in-vivo* experiments (Blasdel, 1992; Bonhoeffer and Grinvald, 1991, 1993; Obermayer and Blasdel, 1993).

**Figure 1:**
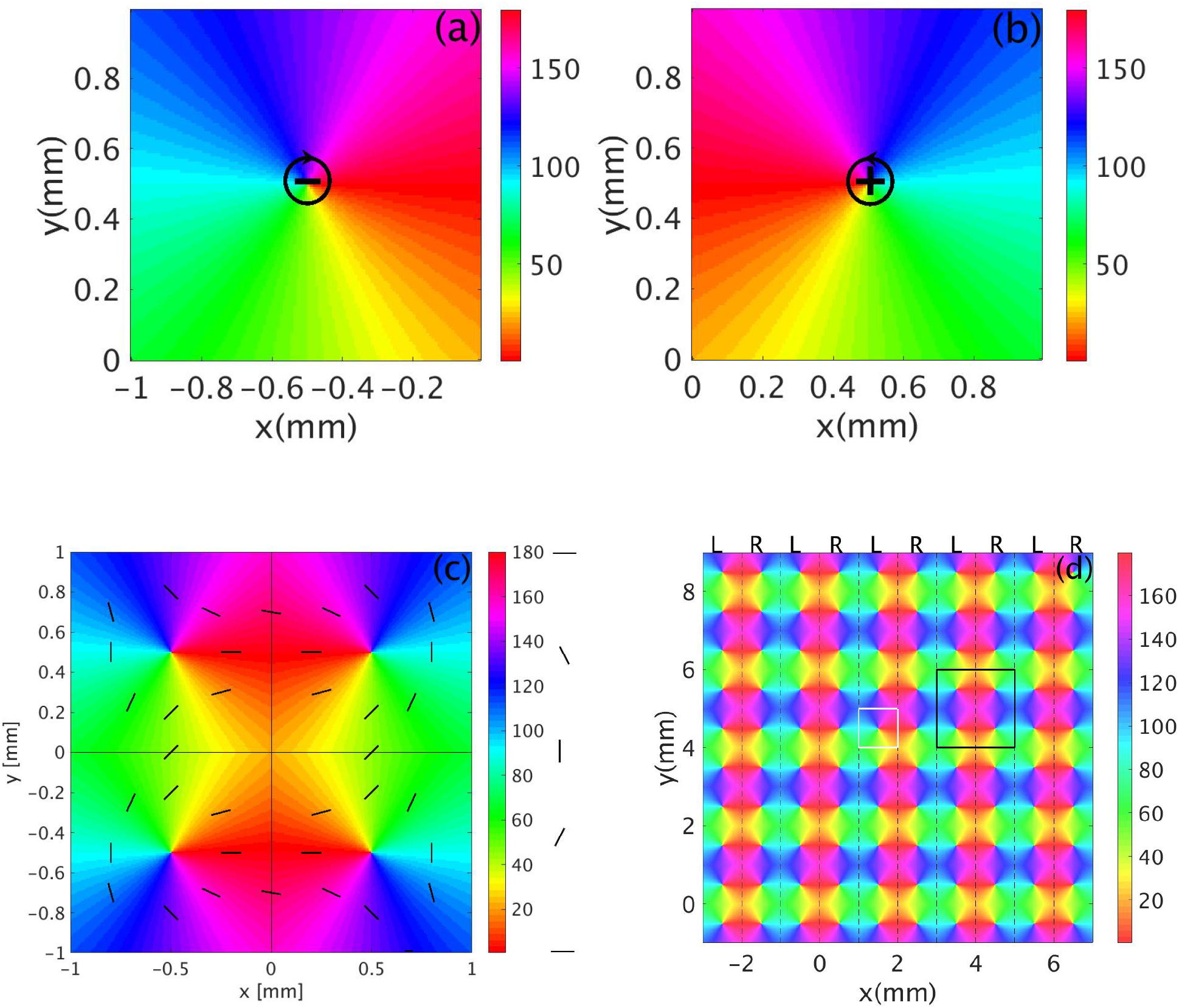
Schematics of visual feature preference maps in V1 with color bars indicating OP in degrees. (a) Negative pinwheel. (b) Positive pinwheel. (c) Lattice unit cell (hypercolumn). The vertical line divides the unit cell into left and right OD columns of equal width, while the horizontal and vertical lines split the unit cell into four squares, each containing one OP pinwheel. The short bars highlight the OP at various locations. (d) Periodic spatial structure of OP and OD columns across a small piece of V1 comprising 25 unit cells. Dashed lines bound left (L) and right (R) OD columns. One pinwheel is outlined in white and one unit cell is outlined in black. Frames (a) and (b) are adapted from Kukjin et al. (2003).

An additional feature of V1 is that regions of similar OP are preferentially linked within and between unit cells by lateral connections, forming a patchy network (Gilbert and Wiesel, 1983; Rockland and Lund, 1982). Furthermore, these patchy connections are concentrated toward an axis that corresponds to the OP of the neurons involved, so the projections from a given unit cell depend strongly on the OP at the source neurons within that cell and are thus strongly anisotropic (Bosking et al., 1997).

When one considers activity on the network, numerous experiments and studies (Eckhorn et al., 1988; König et al., 1995; Singer and Gray, 1995; Engel et al., 1990; Hata et al., 1991; Gray et al., 1989) have shown that neurons with similar feature preference in V1 exhibit synchronized gamma band (30 - 70 Hz) oscillations when the stimulus is optimal, by measuring the multi-unit activities (MUA) and local field potentials (LFP) in area 17 of cats using multi-electrodes. They also showed that the corresponding two-point correlation functions of MUA or LFP commonly have peaks at zero time-lag. Moreover, these synchronized gamma oscillation in V1 arise from the spatial structure of V1, modulated by the specific feature preferences involved. It also has been argued that such synchronized oscillation in gamma band may be involved in visual perception, the binding of related features into unified percepts, and the occurrence of visual hallucinations (Gray et al., 1990; Engel et al., 2001; Bressloff et al., 2002; Siegel et al., 2011; Henke et al., 2014).

Previous theoretical studies (Robinson, 2005, 2006, 2007) used neural field theory (NFT) with patchy propagators to show that patchy connectivity could support gamma oscillations with correlation properties whose features resembled those of some of the experiments noted above. However, the effect of OP in the patchy propagators was not incorporated and the correlations were only explored as functions of one spatial dimension.

In this paper, we generalize and explore the spatiotemporal correlation functions of Robinson (2006, 2007) to two spatial dimensions, and account for the effect of OP on the patchy propagators. We then compare the resulting spatiotemporal correlations with several MUA experiments. In Sec. 2, we briefly describe the relevant aspects of NFT including patchy propagators. Section 3 describes a shape function, which modulates the connection strength between cortical locations that have similar feature preference. In Sec. 4, we derive a general two-point correlation function, and apply this result in Sec. 5 to derive the 2D correlation function in V1 via the linear NFT transfer function of V1. The properties of these correlation functions are explored in Sec. 6, including their predictions for oscillation frequency, time decay, effects of the spatial separation between the measurement points, and the modulation by the OP in V1. The predictions compared with specific experimental outcomes in Sec. 7, and the results are summarized and discussed in Sec. 8.

## 2 Theory

In Sec.2.1 we first review the neural field equations of interest; and, then in Sec. 2.2 we describe the corticothalamic model (Robinson et al., 1997; Robinson, 2005, 2006, 2007), from which we derive a reduced corticothalamic model used in the remainder of this work.

### 2.1 Neural Field Theory

NFT averages neural properties and activity over a linear scale of a few tenths of a millimeter to treat the dynamics on scales larger than this, which is appropriate for the present applications (Deco et al., 2008; Robinson et al., 2005). Each type of neuron is denoted by a subscript *a*. The continuum soma potential *V_a_* is approximated by adding up *V_ab_* resulting from activities of all possible types of synapse on neurons in the spatially extended population *a* from those of type *b*. Thus,

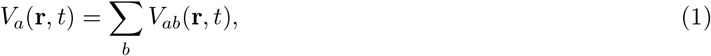

where **r** is the actual spatial location on the cortex, approximated as a two-dimensional sheet, and *t* denotes the time. In the Fourier domain, Eq. (1) can be written as

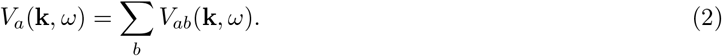

Due to the dependence of *V_ab_* on the synaptic dynamics, signal dispersion in the dendrites, and soma charging, the soma potential corresponding to a delta function input can be approximated by

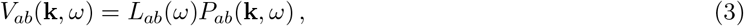

where *P_ab_* is the Fourier representation of the weighted average arrival rate of incoming spikes at each location **r** and time *t*, and *L_ab_* is the function of the synapse-to-soma response in frequency domain,

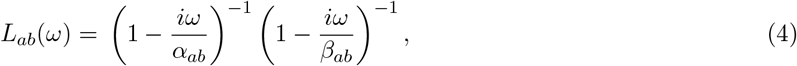

where *α_ab_* and *β_ab_* are the decay and mean rise rate of the soma response.

Action potentials of cells with voltage-gated ion channels are produced when the soma potential exceeds a threshold *θ_a_*. Thus, the mean population firing rate *Q_a_* can be related to the mean soma potential *V_a_* by a nonlinear sigmoid function

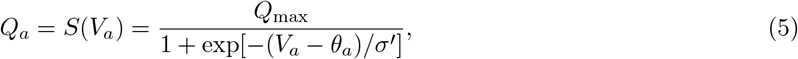

where *Q*_max_ is the maximum possible firing rate, 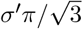 is the standard deviation relative to the mean firing threshold *θ_a_*. We can linearize Eq. (5) by replacing the the sigmoid function *S_a_* by its slope *ρ_a_* at steady state value of *V_a_*, with

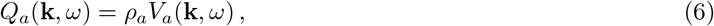

The values of *P_ab_* in Eq. (3) depends on the firing rate *Q_b_* at various source locations and at earlier times (Robinson, 2007). It is governed by the spatiotemporal propagation of pulses traveling within the axons of population *b* to population *a*. By adopting a propagator Γ_*ab*_, *P_ab_* can be expressed as

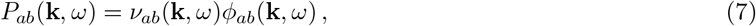

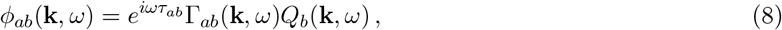

where the forward and inverse Fourier transforms are defined by

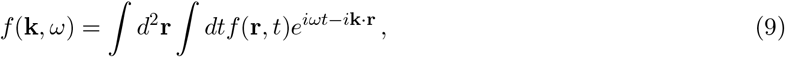

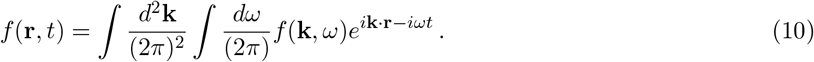

In Eq. (8), *τ_ab_* is the time delay between spatially discrete neuron populations (i.e., not between different **r** on the cortex) and *ν_ab_* represents the coupling between population *b* and population *a*, with

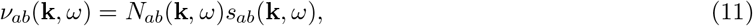

where *N_ab_* is the mean number of synaptic connections to each neuron of type *a* from neurons of type *b* and *s_ab_* is the mean strength of these connections.

Axonal propagation can be approximately described by a damped wave equation (Jirsa and Haken, 1996; Robinson et al., 1997; Schiff et al., 2007)

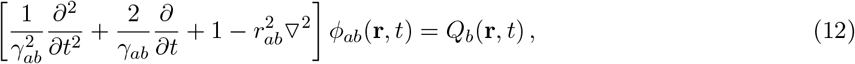

where *γ_ab_* is the temporal damping coefficient and equals *v_ab_*/*r_ab_*, *v_ab_* is the wave velocity, and *r_ab_* is the characteristic range of axons that project from population *b* to *a*.

A uniform-medium propagator 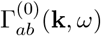 can be derived by first setting the source firing rate *Q_b_* in Eq. (12) to a Dirac delta function of the form: *δ*(**r** − **r′**)*δ*(*t* − *t′*) and replacing *φ_ab_* by 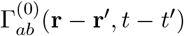. Here, the coordinates (**r**, *t*) represent the spatial location within the target population *a* at time *t*, and (**r′**, *t′*) is the spatial location within the source population *b* at time *t′*. This gives

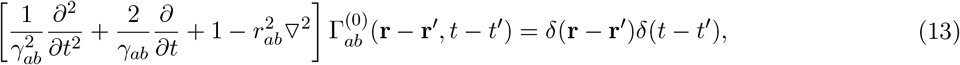

whence

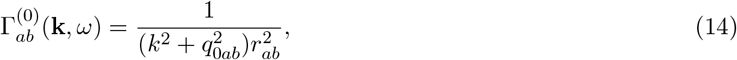

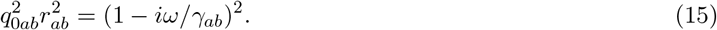

To incorporate the patchy propagation introduced by the periodic cortical structure of V1, periodic spatial modulation is added to the uniform-medium propagator, giving the patchy propagator Γ_*ab*_(**k**, *ω*)

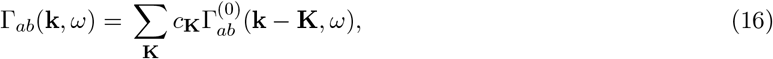

where *c*_**K**_ are the Fourier coefficients of the function that describes the spatial feature preference (i.e., OP), and **K** ranges over the reciprocal lattice vectors of the periodic structure (Robinson, 2007).

In order to perform a linear analysis of the system in later sections, we relate *Q_a_*(**k**, *ω*) and *Q_b_*(**k**, *ω*) via Eqs (1) -(7), which yields

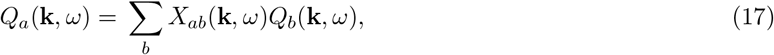

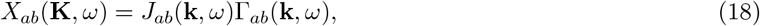

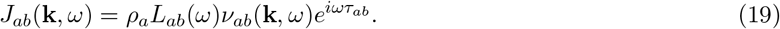

### 2.2 EMIRS corticothalamic model

The previously developed corticothalamic model (Robinson, 2005) consists of five neural populations, which are the long-range excitatory (*e*), midrange excitatory (*m*), short-range inhibitory (*i*), thalamic reticular (*r*), and relay thalamic (*s*) populations; hence, it is termed the EMIRS model. Figure 2(a) shows the full EMIRS model and its connectivities between neural populations. There is only one external input *ϕ_sn_*, which is incident on the relay nuclei.

**Figure 2:**
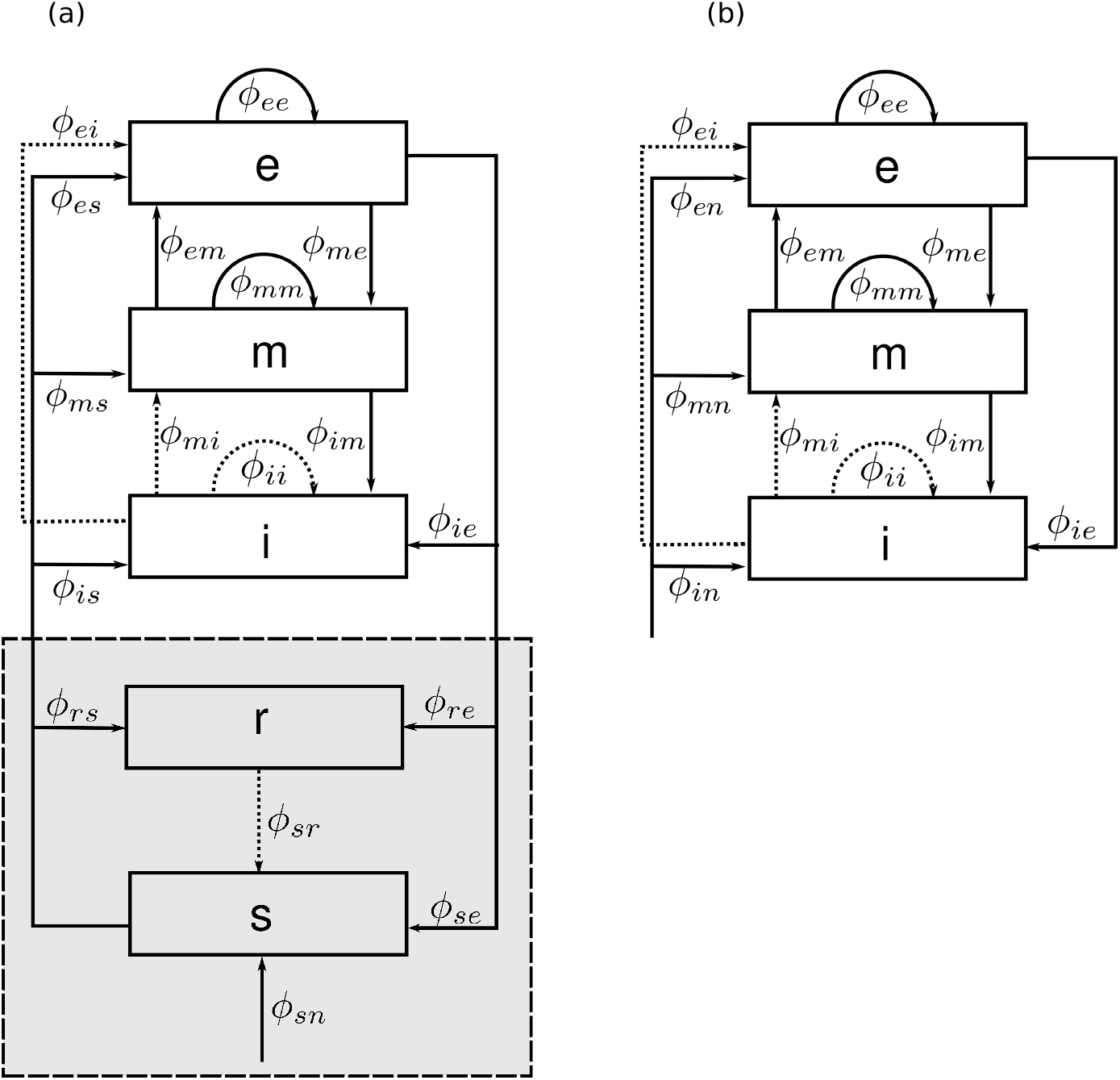
Schematics of the corticothalamic system. (a) The full EMIRS model with the thalamus part shown in the gray rectangle, each *ϕ_ab_* quantifies the connection to population a from population b. (b) The simplified EMIRS model with the thalamic part approximated as a cortical input.

In this work, we are mainly concerned with cortical neural activities in the gamma band (30Hz - 70Hz), which are higher than the resonant frequency (~10Hz) of the corticothalamic loops. This enables us to neglect the corticothalamic feedback loops of the full EMIRS model, leading to the reduced model in Fig. 2(b). This model only includes the cortical excitatory, mid-range, and short-range inhibitory populations, and the signals from the thalamus are treated as the input to the cortex. Thus, rather than having feedback inputs from the thalamus, we approximate these inputs as a common external input *ϕ_an_* to the cortex. The subscript *a* denotes the three cortical neural populations (*e, m, i*).

## 3 Patchy Propagation

Patches of neurons with similar feature preference in the visual cortex are preferentially connected (Bauer et al., 2014; Bressloff and Cowan, 2003; Gilbert and Wiesel, 1983; Lund et al., 2003; Muir et al., 2011). Moreover, these patchy projections are not spread randomly, but are concentrated toward an axis corresponding to the OP angle of the neurons (Bosking et al., 1997; Malach et al., 1993; Sincich and Blasdel, 2001). To model this overall modulation of the propagation use an elliptic Gaussian shape function whose long axis is oriented at the local OP *ϕ* at the source point **r′** of outgoing axons. If **r′** = (*x′*, *y′*) and **r** = (*x, y*), we have

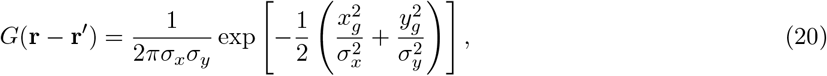

where

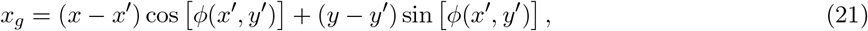

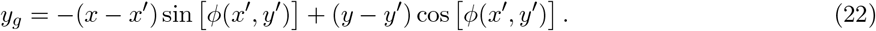

Here *σ_x_* = 2.6 mm and *σ_y_* = 0.7 mm are the spatial ranges along the preferred *x_g_* and orthogonal *y_g_* directions, with values chosen to match the experiment findings in tree shrew by Bosking et al. (1997).

Figures 3(a) and (b) show contour plots of *G*(**r** − **r′**) for OPs of 0° and 45°, respectively and source points **r′** within a central unit cell (see Fig. 1(c)), which is outlined by the red square. The gray scale shows the strength and direction of spatial modulation of the propagation. We see that the propagation of activity at **r′** is modulated in such a way that it propagates further along the axis of preferred orientation, and it propagates less along the axis of orthogonal orientation Bosking et al. (1997).

**Figure 3:**
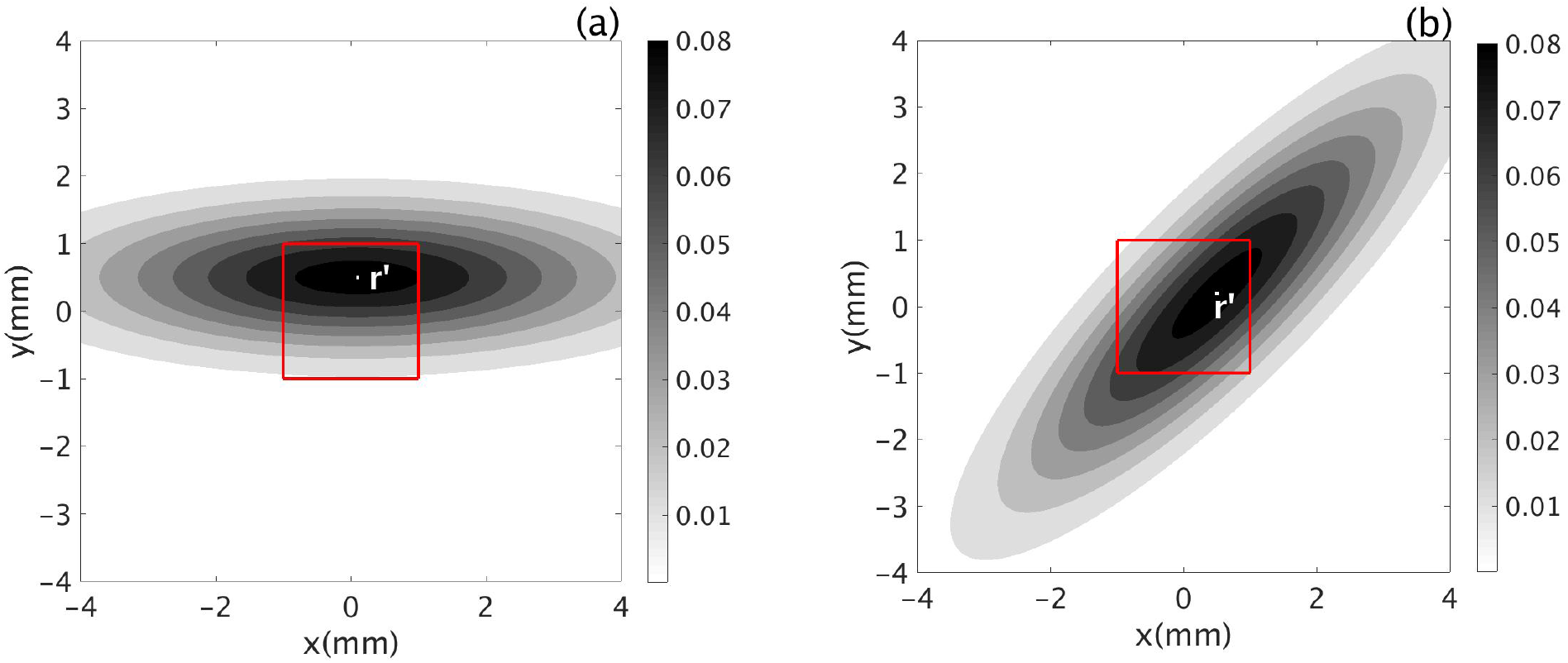
Plots of Eq. (20) with the central unit cell outlined in red; the color bar shows values of *G*(**r** − **r′**). (a) OP = 0°. (b) OP = 45°.

Patchy propagation also has modulation with spatial period *k* = 2*π*/*a* in both preferred and orthogonal directions, where *a* ≈ 2 mm is the width of the unit cell. To incorporate this modulation, we multiply the oriented elliptic Gaussian function by an isotropic cosine function. The final form of the shape function is then

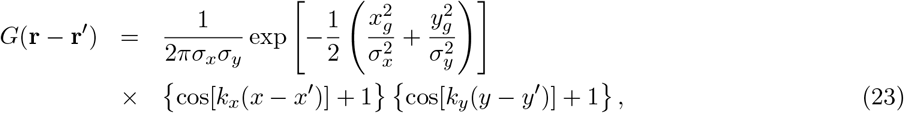

where *k_x_* = *k_y_* = 2*π*/*a*.

Figures 4 (a) and (b) show the resulting contours of Eq. (23), for *ϕ*(**r′**) = 0° and 45°, with *σ_x_* = 2.6 mm and *σ_y_* = 0.7 mm. For both cases, when **r** — **r′** < 0.5 mm the underlying neurons respond to the stimulus, regardless of their orientation preference. In the case of ϕ(**r′**) = 45°, we can see regions of contours with weaker connection strength, which represent patches of the cortex that have the same OP angle as the source point but are located slightly off the main propagation axis.

**Figure 4:**
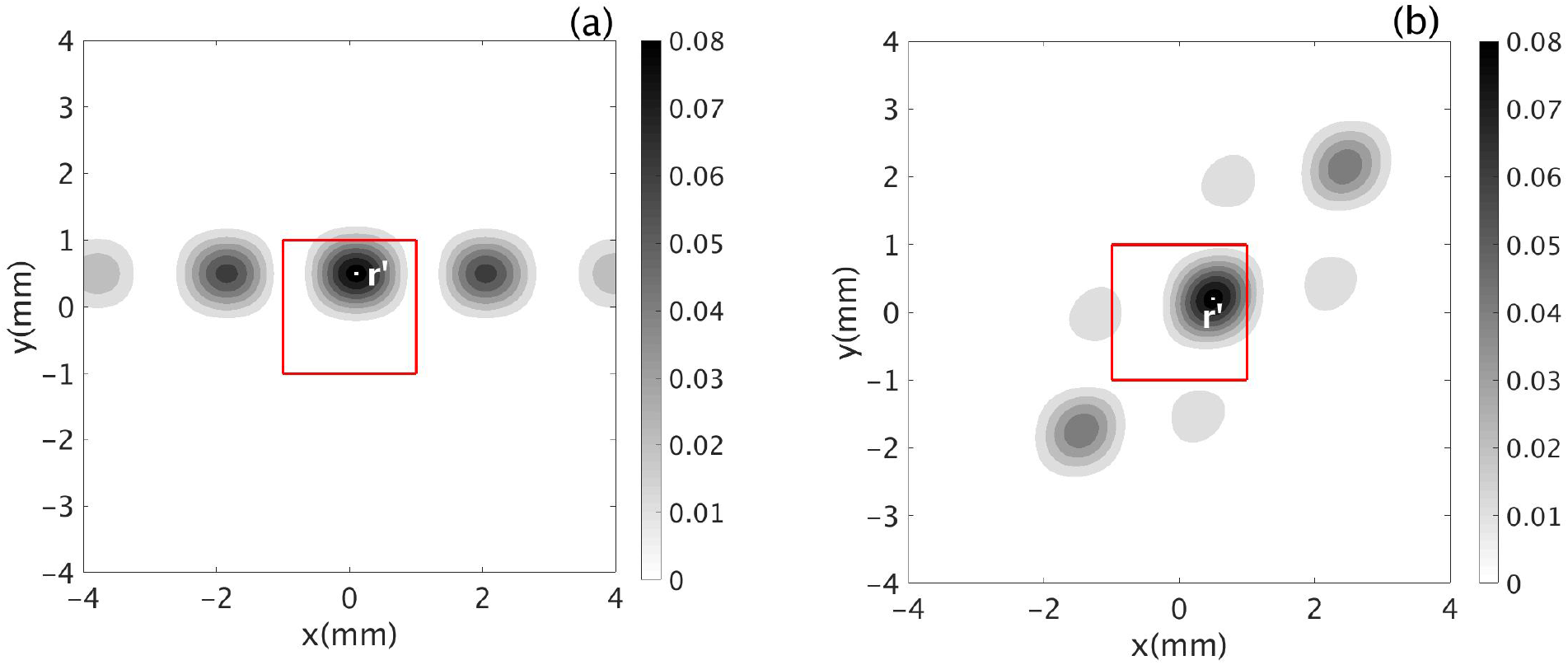
Contour plot of the shape factor *G*(**r**, **r′**) in Eq. (23) with the central unit cell outlined in red and containing the source point **r′**. The color bar shows the values of *G*(**r**, **r′**). (a) *ϕ*(**r′**) = 0°. (b) *ϕ*(**r′**) = 45°.

## 4 General Correlation Function

This section summarizes the derivation of the general two-point correlation function between the cortical firing rates measured at two different locations, generalizing the analysis of Robinson (2007) and improving its notation.

We first assume the visual cortex receives two uncorrelated and spatially localized inputs (stimuli) at locations **s**_1_ **s**_2_. Further, cortical activity is measured at locations **m**_1_, **m**_2_. Figure 5 shows a schematic of typical spatial locations and OPs involved in required to deriving the correlation function. The two ellipses in solid green and red lines, centered at **s**_1_ and **s**_2_ represent the shape factor *G*(**r** − **r′**) for OPs *ϕ*(**s**_1_) = 45° and *ϕ*(**s**_2_) = 0°. The arrows indicate propagation of neural activity from sources **s**_j_ to measurement points **m**_*k*_.

We first derive equations for the neural activities at **m**_1_ and **m**_2_ due to inputs at **s**_1_ and **s**_2_. The activity **Φ**_1_ at **m**_1_ can be written as

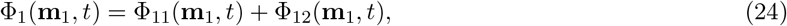

where **Φ**_11_(**m**_1_, *t*) denotes the activity propagated to **m**_1_ from **s**_1_ and **Φ**_12_ (**m**_1_, *t*) denotes the activity propagated to **m**_1_ from **s**_2_. Similarly, activity at **m**_2_ is

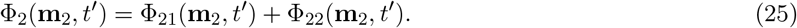

We can write **Φ**_11_ as

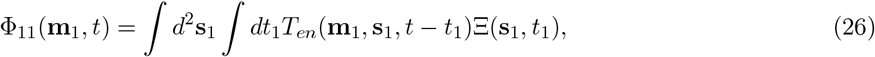

where *T_en_* (**m**_1_, **s**_1_, *t* − *t*_1_) is the transfer function that relates the activities at **m**_1_ to the stimulus Ξ at **s**_1_. From Robinson (2007), Ξ(**s**_1_, *ω*) can be approximated in the spatial and Fourier domains as

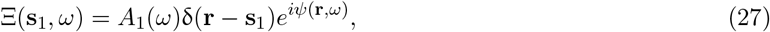

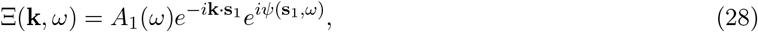

where the real quantities *A*_1_(*ω*) and *ψ*(**s**_1_,t_1_) are the amplitude and the phase of the input at **s**_1_. We Fourier transform Eq. (26) and use Eq. (28) to get

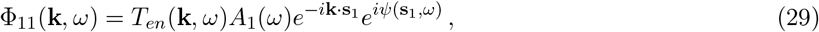

Likewise,

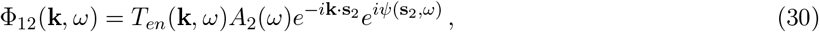

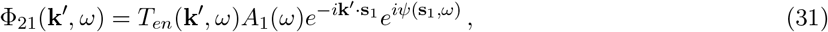

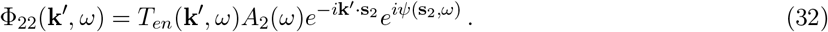

Combining Eqs (29) - (32), we have

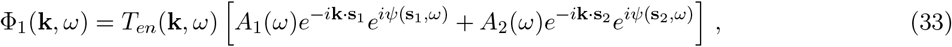

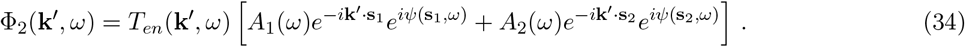

The general two-point correlation function at measurement locations **m**_1_ and **m**_2_ is (Robinson, 2007)

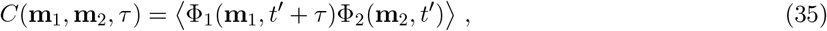

where *t* = *t* − *t′*, and the angle brackets refer to the averages over *t′* and over the phase of the inputs. We use continuous Fourier transforms and integration over *t′* to achieve the averaging, which yields,

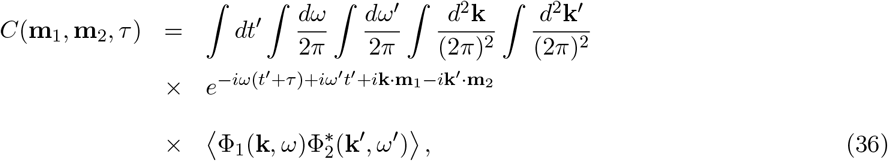

Evaluating the integrals with respect to t’ and ω’ yields

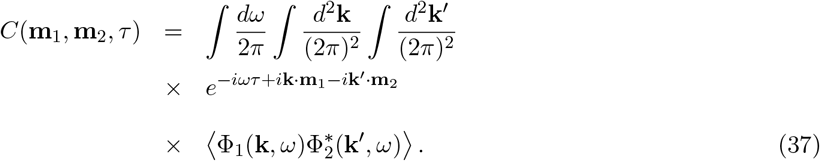

Substituting Eqs (33) and (34) into Eq. (37), and taking the inverse Fourier transform then gives

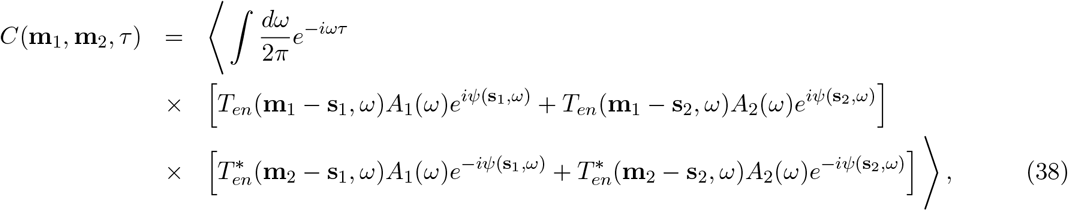

where the angle brackets now denote the average over the phases at **s**_1_ and **s**_2_. If the phases of the inputs are random and uncorrelated, we have

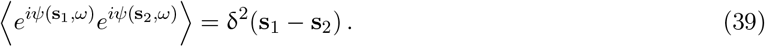

Then the cross terms between **s**_1_ and **s**_2_ in Eq. (38) are zero and one finds

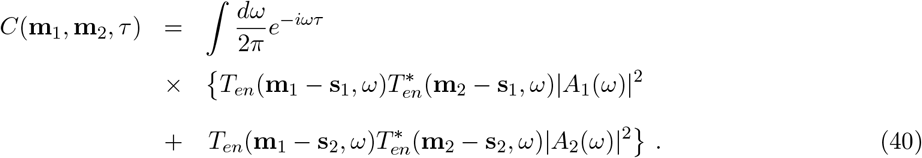

## 5 Gamma oscillations and correlations in the EMIRS Model

In this section we first analyze the frequency responses of the reduced EMIRS model (Sec. 5.1), and then derive the transfer function *T_en_* which allow us to obtain a specific form of the 2D two-point correlation function (Sec. 5.2). Lastly, we study the general properties of the correlation function for several configurations of source and measurement points (Sec. 6).

### 5.1 Resonances of the EMIRS Model

In this subsection, we briefly outline the derivation of the linear transfer function of the EMIRS model and its resonances Robinson (2006, 2007).

When only the cortical populations of the model are considered, the linear transfer function *T_en_*(**k**, *ω*) can be written as

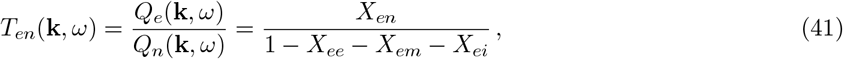

where *X_ab_* is defined in Eqs (4), (18), and (19).

The resonances of the system arise from the poles of the transfer function, which are given by setting the denominator of Eq. (41) to zero. For *k* ≫ 1/*r_ee_*, |*X_ee_*| ≪ |*X_ei_*| so *X_ee_* can be neglected and the resonance condition becomes

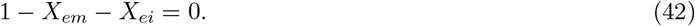

Substituting Eqs (4), (14), (18), and (19) into Eq. (42) gives

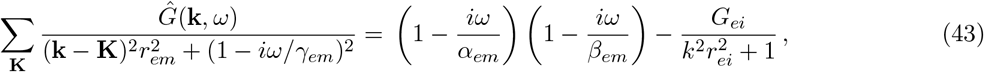

where

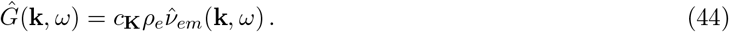

When **k** ≈ **K**, the denominator on the left hand side of Eq. (43) is small, and the corresponding term dominates the sum over the lattice vectors **K**. Assuming *Ĝ*(**k**, *ω*) is purely spatial, *Ĝ*(**k**, *ω*) can be written as *Ĝ*(**K**) and Eq. (43) becomes

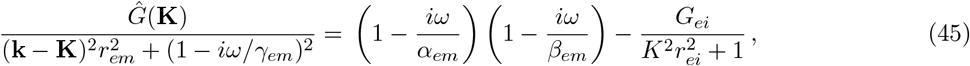

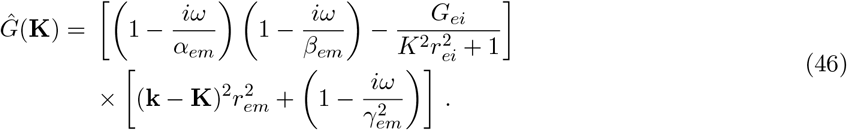

Expanding Eq. (46) by multiplying each term out, letting 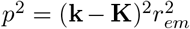, and omitting subscripts *em* we find

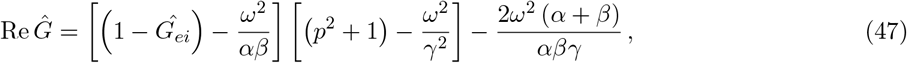

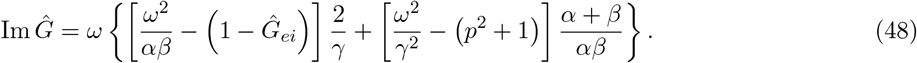

Real *Ĝ* are considered here for a simpler case of direct response from the stimuli, and for ignoring the phases shift. If *Ĝ* is positive, the resonance occurs only at *ω* = 0 and it is irrelevant to gamma band oscillation. So we choose negative *Ĝ*, and the exact resonance occurs at Ω given by

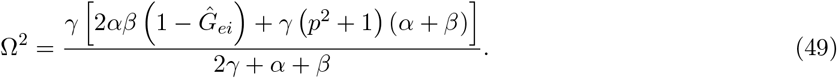

### 5.2 Derivation of the corticothalamic transfer function in closed form

Robinson (2007) approximated the transfer function using only the lowest reciprocal lattice vector K as

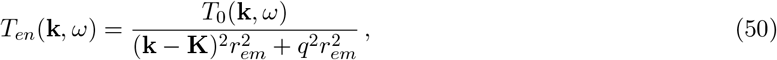

where

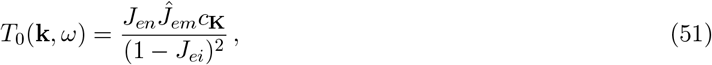

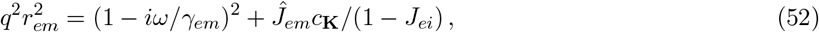

where *Ĵ_em_* is defined in Eq. (19).

We now include higher order lattice vectors to incorporate the finer spatial structure of the OP map into our analysis. From the analysis in Sec. 5.1, we know that sharp resonances of the system can only occur at its frequency pole pairs at (±**K**_*j*_, ±Ω_*j*_), where **K**_*j*_ = ±**K**_1_, ±**K**_2_, ±**K**_3_, ⋯, are the reciprocal lattice vectors and Ω_*j*_ = ±Ω_1_, ±Ω_2_, ±Ω_3_, ⋯, are the resonance frequencies corresponding to each **K**_*j*_. Let **k** ≈ **K**_*j*_ and *ω* ≈ Ω_*j*_ for *T*_0_, Eq. (50) can be approximated as the sum of the transfer functions of each frequency pole pair:

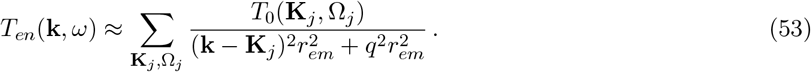

Spatially Fourier transforming Eq. (53) gives

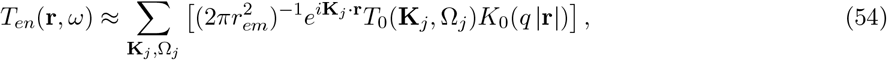

where *T*_0_ and *q* are given in Eqs (51) and (52), and *K*_0_ is a modified Bessel function of the second kind (Olver et al., 2010). Substituting Eq. (54) into Eq. (40) and letting |*A*_1_(*ω*)| = |*A*_2_(*ω*)| = 1 for simplicity, gives us the final equation for the EMIRS spatiotemporal correlation function

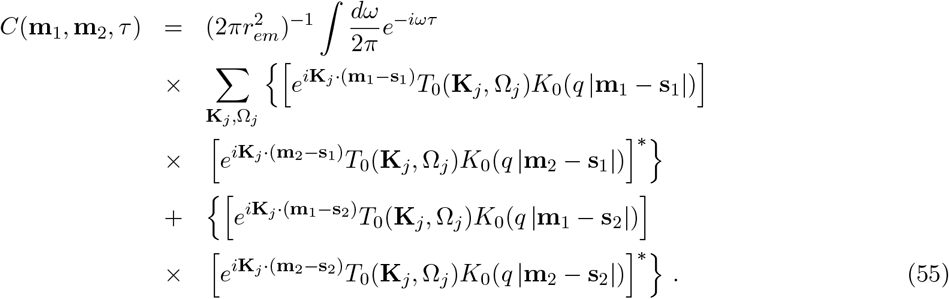

Some general aspects of Eq. (55) are that the correlations fall off on a characteristic spatial scale of (Re*q*)^−1^ because *K*_0_(*z*) ~ exp(—*z*) at large *z* in the right half plane. For the same reason, there is an oscillation with spatial frequency of Im*q*. As we see below, resonances in *T*_0_ select dominant temporal frequencies in the correlations.

## 6 Spatiotemporal properties of the correlation function

Here, we first explore the temporal properties of the correlation function predicted using Eq. (55). Then we explore its spatial properties with a single input. Lastly, we examine the spatial correlation in the case of two input sources.

In all the cases described below, the correlation is calculated by numerically evaluating Eq. (55) and locating **m**_1_, **m**_2_, **s**_1_, and **s**_2_ under different conditions. These conditions include using different optimal OPs for the measurement points and source points, and varying the distances between the measurement points. The results are presented in Fig. 6. All correlations are normalized such that *C*(**m**_1_, **m**_2_, *τ*) = 1 when **s**_1_ = **s**_2_, and **m**_1_ = **m**_2_ are placed very close to the sources. Table 1 summarizes the parameters we use for the calculations.

**Figure 5:**
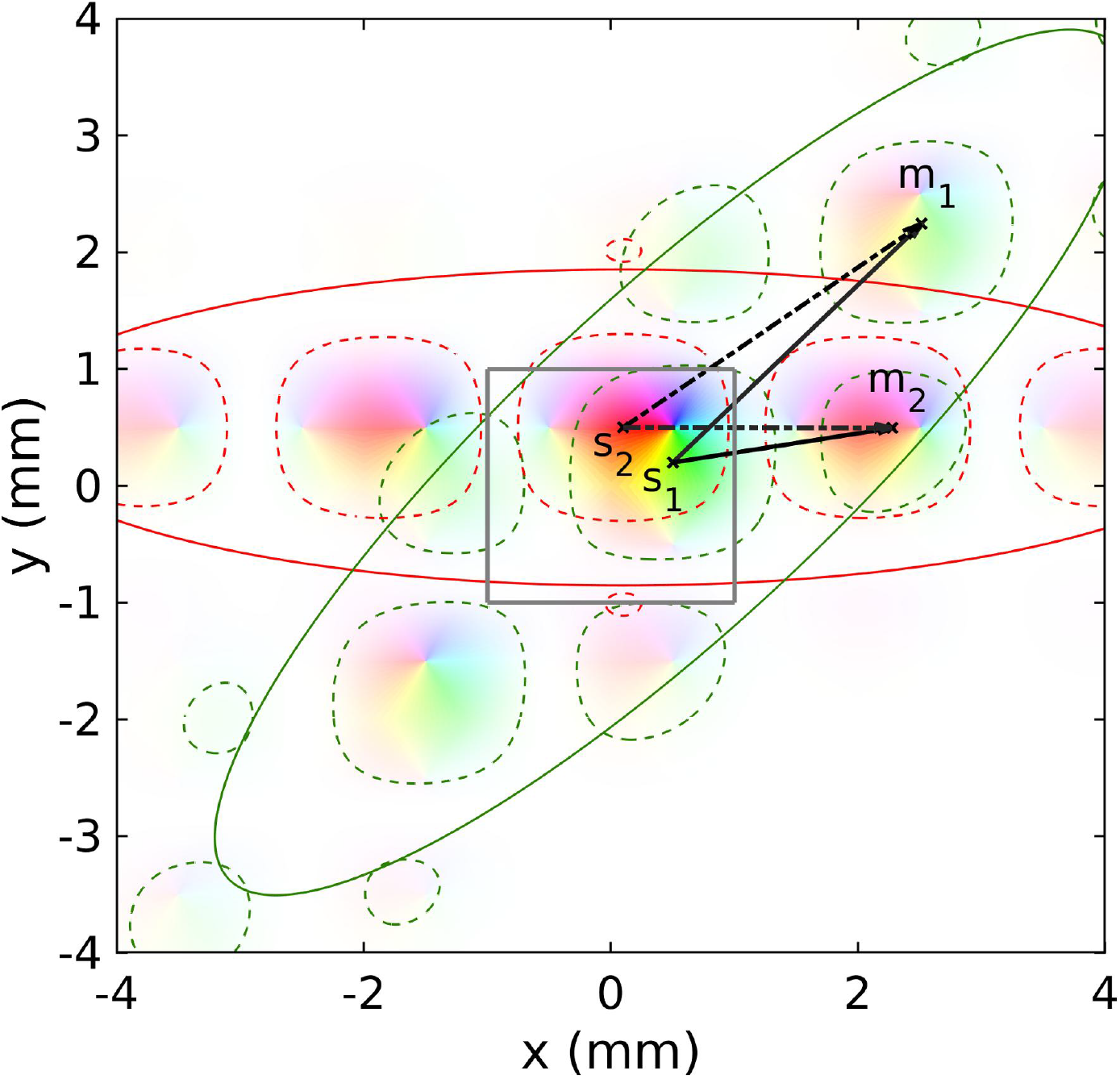
Schematic for deriving the correlation functions, where **s**_1_ and **s**_2_ denote the stimulus/source points and **m**_1_ and **m**_2_ are the measurement points. The ellipses in solid green (red) lines indicate the overall shape of the orientation-modulated propagation from **s**_1_ (**s**_2_), as given by Eq. (23). The ellipsoids outlined in dotted green (red) lines indicate the patchiness of the propagation along the OP of **s**_1_ (**s**_2_), with period *k* = 2*π*/*a*. The solid and dash-dotted arrows denote propagation from **s**_1_ and **s**_2_, respectively, to **m**_1_ and **m**_2_.

**Table 1:**
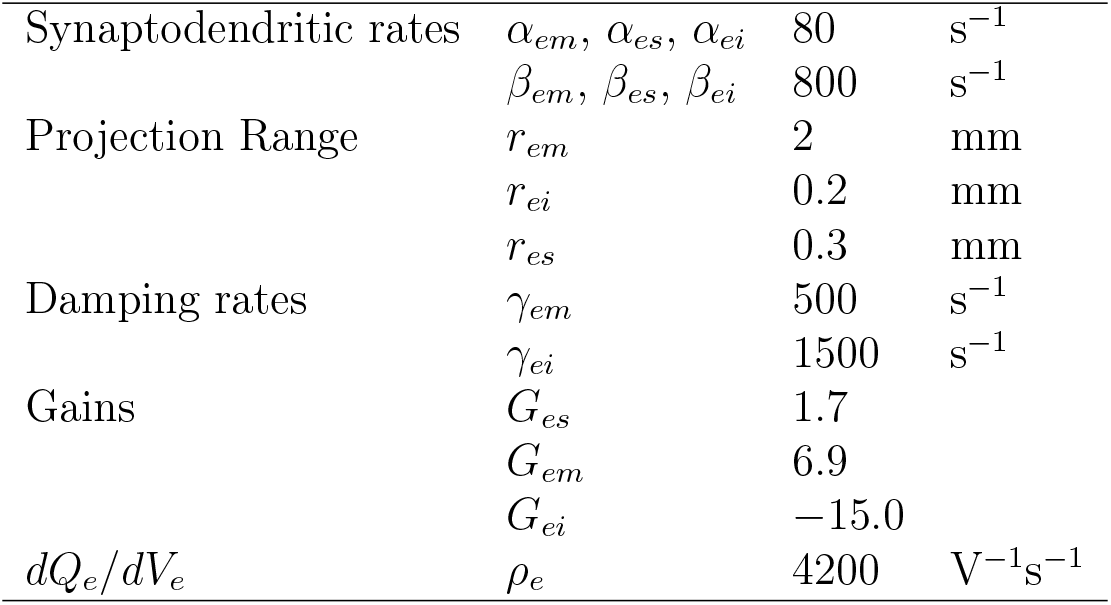
EMIRS model parameters

### 6.1 Temporal Correlation Properties

In Fig. 6(a), we illustrate the temporal correlations evoked by binocular stimulation when s1 and s2 have the same OP *ϕ*(**s**) = 90° and **s**_1_ and **s**_2_ are located in the same unit cell, but in different OD columns. The strength of propagation of neural signals from two sources is indicated by contour lines of Eq. (23); the propagation is predominantly parallel in this case.

**Figure 6:**
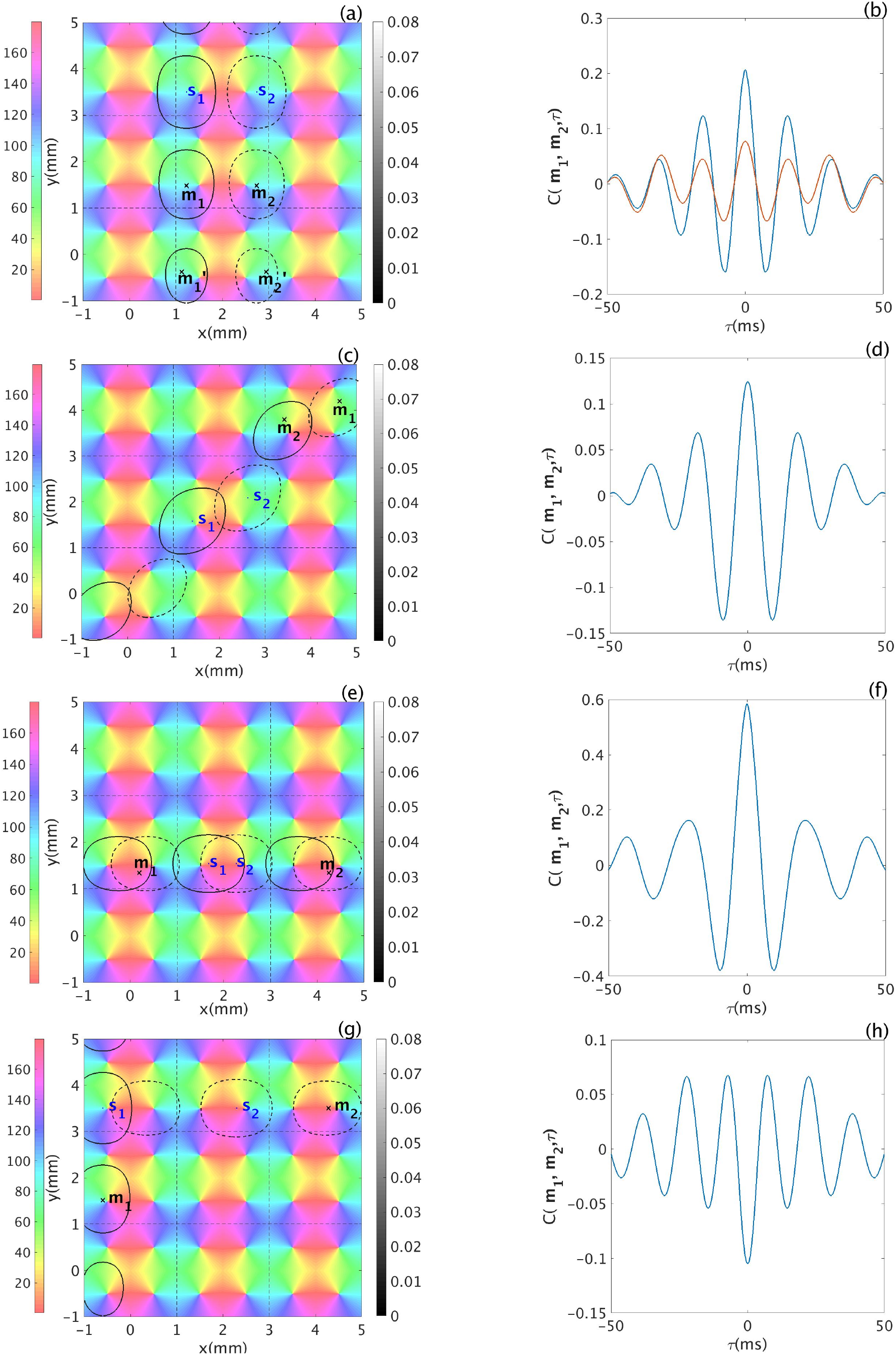
Properties of temporal correlations. (a) Locations of measurement points **m**_1_ and **m**_2_, and source points **s**_1_, and **s**_2_ within 9 unit cells in V1. All points have OP of 90°. The color bar shows the OP and the gray contours show the strength of propagators given by Eq. (23) with solid and dashed curves for propagation from **s**_1_ and **s**_2_, respectively. (b) Temporal correlations. Blue curve shows *C*(**m**_1_, **m**_2_, *τ*), while the orange curve shows 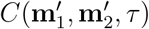. (c) As for (a) but with all points have OP of 45°. (d) Temporal correlation for (c). (e) As for (a) but with all points have OP of 0°. (f) Temporal correlation for (e). (g) As for (a) but with OP of the sources are orthogonal. Furthermore, OP at **s**_1_ is optimal for **m**_1_ whereas OP at **s**_2_ is optimal to **m**_2_. (h) Temporal correlation for (g).

The points **m**_1_ and **m**_2_ are located in a different unit cell to the source points; are approximately 2 mm away from each other; and are located at approximately 2 mm from their respective collinear source points, the OPs at **m**_1_ and **m**_2_ are also 90°. We have also placed additional measurement points 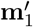 and 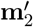, with the same OP as **m**_1_ and **m**_2_, but are approximately 4mm from the sources.

Figure 6(b) shows the temporal correlation functions *C*(**m**_1_,**m**_2_, *τ*) and 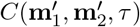. Both oscillate at around 64 Hz, in the gamma range. Furthermore, each has a peak centered at *τ* = 0, so the neural activities at **m**_1_ and **m**_2_, 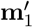 and 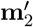, are synchronized. The time for their envelopes to decrease to 1/*e* (~ 30%) of the peak value is ≈ 18 ms. However, when the measurement points are placed further away from the sources, the correlation at *τ* = 0 becomes weaker, as seen by comparing the two curves.

Figure 6(c) shows a case for which the OP of all sources and measurement points is equal (at 45°). Figure 6(d) shows that the resulting correlation also has a central peak at zero time-lag, oscillates in the gamma band at ~55 Hz, and its envelope decreases by 1/*e* at *τ* ≈ 21 ms.

In order to explore the correlation properties between OD columns, we place all the source points and measurement points co-linearly with OP = 0° in Figure 6(e). Synchronized activities at **m**_1_ and **m**_2_ are shown by the center peak at *τ* = 0 in Fig. 6(f). This correlation also exhibits gamma band oscillation at ≈ 50Hz, and the decrease by 1/*e* from the peak happens at ≈ 21ms. One thing worth to be mentioned here is that the correlation strength due to inter-columnar connection shown in Fig. 6(f), is stronger than the intra-column connection in Fig. 6(b).

To further investigate the correlation properties, Fig. 6(g) shows a case in which the two measurement sites have orthogonal OPs, and so does the sources: the OP at **s**_1_ and **m**_1_ is 90°, while at **s**_2_ and **m**_2_ it is 0°. The distance between the two measurement points is around 5.5 mm. In this case, **s**_1_ tends to evoke strong response at **m**_1_, but not at **m**_2_. This introduces an anticorrelation between **m**_1_ and **m**_2_. Similarly, adding another source s2 only stimulates **m**_2_ and it again makes the activities at two measurement sites anticorrelated. This negative correlation is exactly shown by our predicted result in Fig. 6(g). It displays a negative peak at *τ* = 0.

### 6.2 Two dimensional correlations due to a single source

To demonstrate how the correlation strength is influenced by the location of the measurement sites and their OP, we fix the location of a source **s**_1_ and a measurement point **m**_1_, as in Fig. 6(a). We then map the correlation with the second measurement point **m**_2_ at *τ* = 0 as a function of the latter’s position on V1. The resulting map is shown in Fig. 7, normalized to the maximum value of *C*(**m**_1_, **m**_2_, 0).

**Figure 7:**
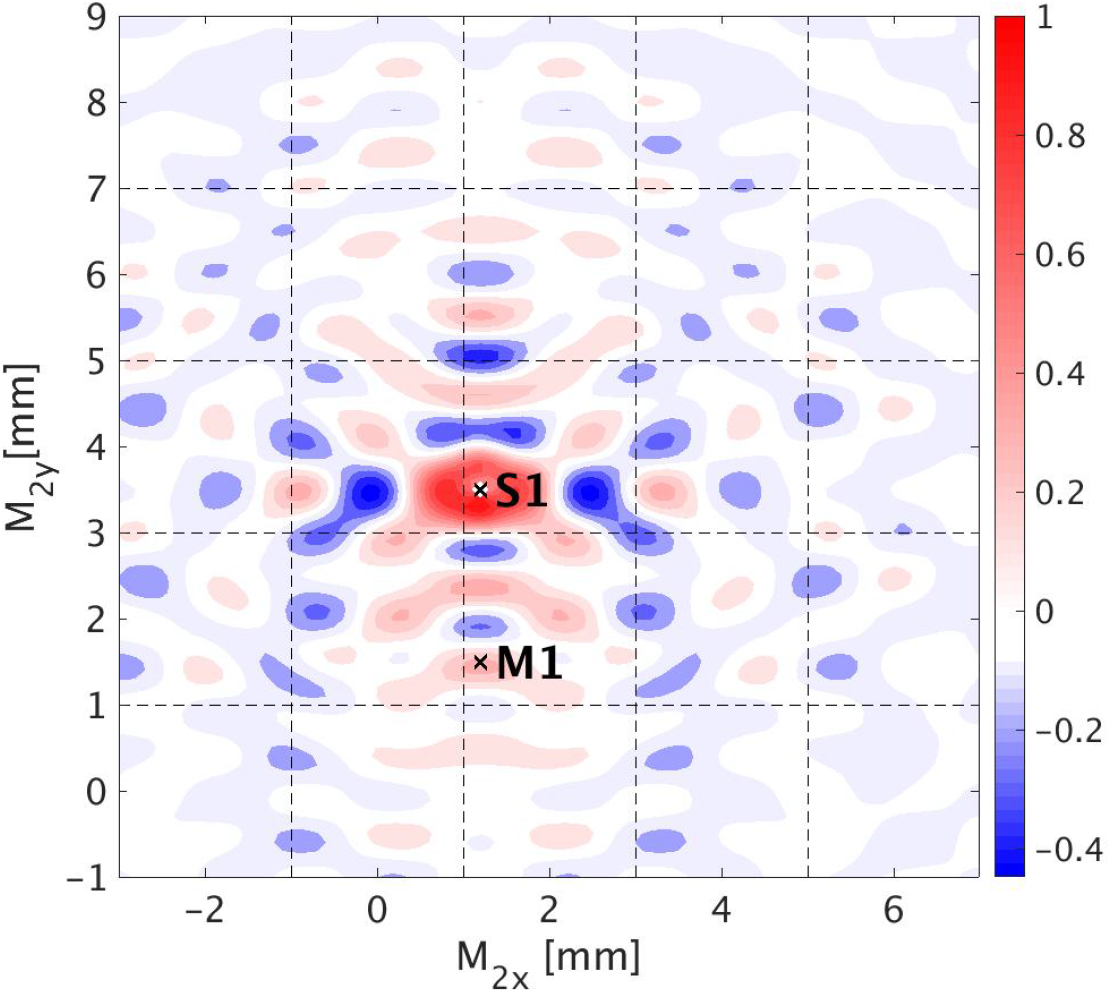
Normalized contour plot of *C*(**m**_1_, **m**_2_, 0) on V1, from Eq. (55) with a single input at **s**_1_. The locations of **s**_1_, and measurement point **m**_1_ are fixed and the location of measurement point **m**_2_(*x,y*) is given by the axes. The location and OP of **s**_1_ and **m**_1_ are the same as shown in Figure 6(a). The color bar indicates the strength of the correlation. Dashed lines bound unit cells.

Figure 7 shows that: (i) The strongest positive correlations are located along a vertical axis passing through the source point **s**_1_ whose OP is 90°; (ii) Patterns of the correlated regions are almost symmetric around the vertical axis in (i); (iii) The correlation strength falls off with distance between the two measurement points, as expected from Eq. (23); In addition, the correlation nearly vanishes when the measurement sites are greater than 7 mm apart, and this agrees with the experimental results, which suggested that oscillatory cross-correlations are not observed when the spatial separation of neurons exceeds 7mm. (iv) The central peak shows that when the distance between **m**_2_ and **s**_1_ is less than 0.5 mm, the correlations are strong and do not depend on the OPs at these locations, in accord with experiments (Bosking et al., 1997; Engel et al., 1990; Gray et al., 1989; Swindale, 1996). (v) The positive correlations correspond to regions of OP approximately equal to **s**_1_’s OP. while negative correlation regions correspond to OPs approximately perpendicular to the source OP angle. This shows that only neurons with similar OP to the source respond to the input stimulus.

### 6.3 Two dimensional correlations due to two sources

Here we explore the dependence of the correlation function *C*(**m**_1_, **m**_2_,0) on the position of measurement point **m**_2_ with two inputs **s**_1_ and **s**_2_. The location of the measurement points and source points are set up exactly as in the previous case and the additional source **s**_2_ has the same OP as **s**_1_ (i.e. 90°).

The resulting map is shown in Fig. 8 and has similar properties to the previous case with one input, namely, the strongest correlations between the measurements points are along a vertical axis, which matches the OP of the sources. The positive correlation regions along this axis have a spatial period of 1 mm, corresponding to the minimum distance between regions having the same OP angle as the sources. However, the negative correlation regions now tend to align horizontally, which represents the direction orthogonal to the OP. The input source **s**_2_ is not surrounded by positive correlation regions as **s**_1_ is; rather, the negative correlations right above **s**_2_ correspond to a region where the OP of **m**_2_ is ~ 0°. This is consistent with Sec. 6.1, where we showed that measurement points with orthogonal OPs tend to be anticorrelated at *τ* = 0. In that case, we have predicted that when the OP of two measurement points are 0° and 90° respectively, the source that is optimal to one of the measurement site introduces negative correlation between the two.

**Figure 8:**
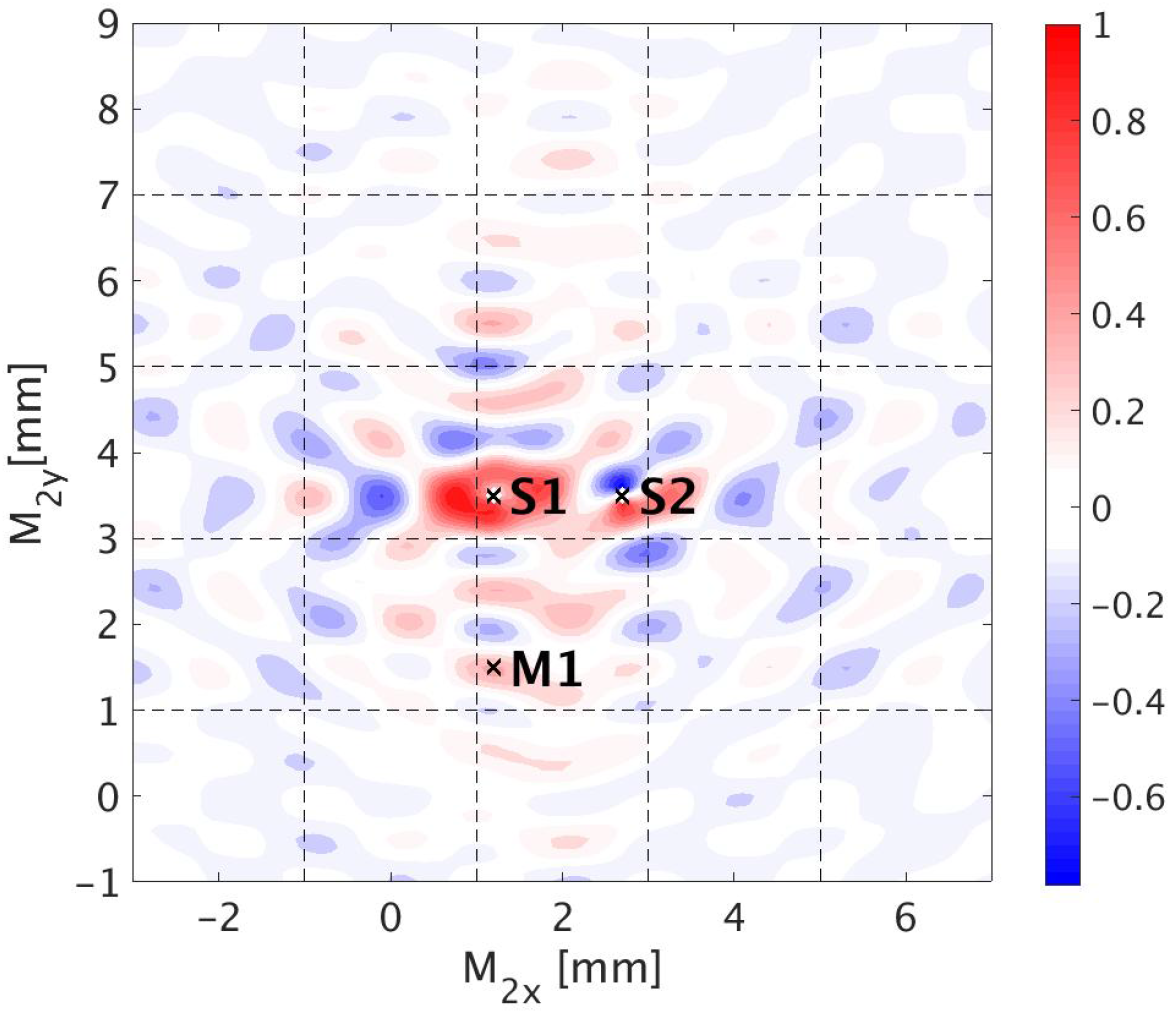
Normalized contour plot of C(**m**_1_, **m**_2_,0) on V1, from Eq.(55) with two inputs **s**_1_ and **s**_2_. The locations of the sources and measurement points **m**_1_ are fixed and the location of measurement point **m**_2_(*x,y*) is given by the axes. The location and OP of **s**_1_, **s**_2_, and **m**_1_ are the same as shown in Figure 6(a). The color bar indicates the strength of the correlation function. Dashed lines bound unit cells.

## 7 Comparison Between Theory and Experiment

In this section, we compare the predicted correlation functions with experimental correlations obtained from Engel et al. (1990), who published temporal correlation functions of MUA and LFP data under various conditions.

### 7.1 Description of the Experiments

In these experiments, the MUA and LFP measurements were recorded from an array of electrodes that were inserted in 5 to 7 spatially separated sites in area 17 of anesthetized adult cats, with neighboring recording sites spaced 400 - 500 *μ*m apart. Oriented light bars were used as binocular stimulation. Each trial lasted for 10 seconds and one trial set was composed of 10 trials with identical stimuli. During each trial, the light bars were projected onto a screen that was placed 1.10 m in front of the eye-plane of the cat. The autocorrelation function (ACF) and cross-correlation function (CCF) of the MUA data were computed. CCFs were calculated on each individual trials first, then averaged to get the final single CCF corresponding to a specific input stimulus Engel et al. (1990).

### 7.2 Mapping experimental conditions to a regular lattice

The experimental stimulation was binocular, so a single moving light bar at a specific point in time, maps to two source points on V1 (**s**_1_ and **s**_2_), both with OP equal to the bar orientation, one located in left OD column and one in the right OD column.

In Engel *et al*.’s experiments, there are five fixed measurement points labeled as **m**_1_ to **m**_5_. Cells at measurement points **m**_1_, **m**_3_, and **m**_5_ have similar orientation preference and are nearly orthogonal to the OP preference of cells at measurement points **m**_2_, **m**_4_. We map these points onto the regular grid used in our model, which results in slight distortion (< 0.5 mm) of the original cortical surface in order to preserve the measurement-point OPs. The OPs of **m**_1_ to **m**_5_, computed after mapping onto our regular lattice match to the OPs given by the experiments within 1°.

Here, we calculate the temporal correlation functions for two sets of experimental conditions, where the only difference between the two is the OP of the stimulus. One stimulus is oriented at 157° and another one oriented at 90°. Figure 9 shows both the stimulation and measurements sites on the idealized OP map. The sources **s**_1_ and **s**_2_ indicate the 157° stimulus, while **s**_3_ and **s**_4_ represent the the 90° stimulus. The locations of the measurement sites **m**_1_ to **m**_5_ are the same for the two sets of experimental conditions.

**Figure 9:**
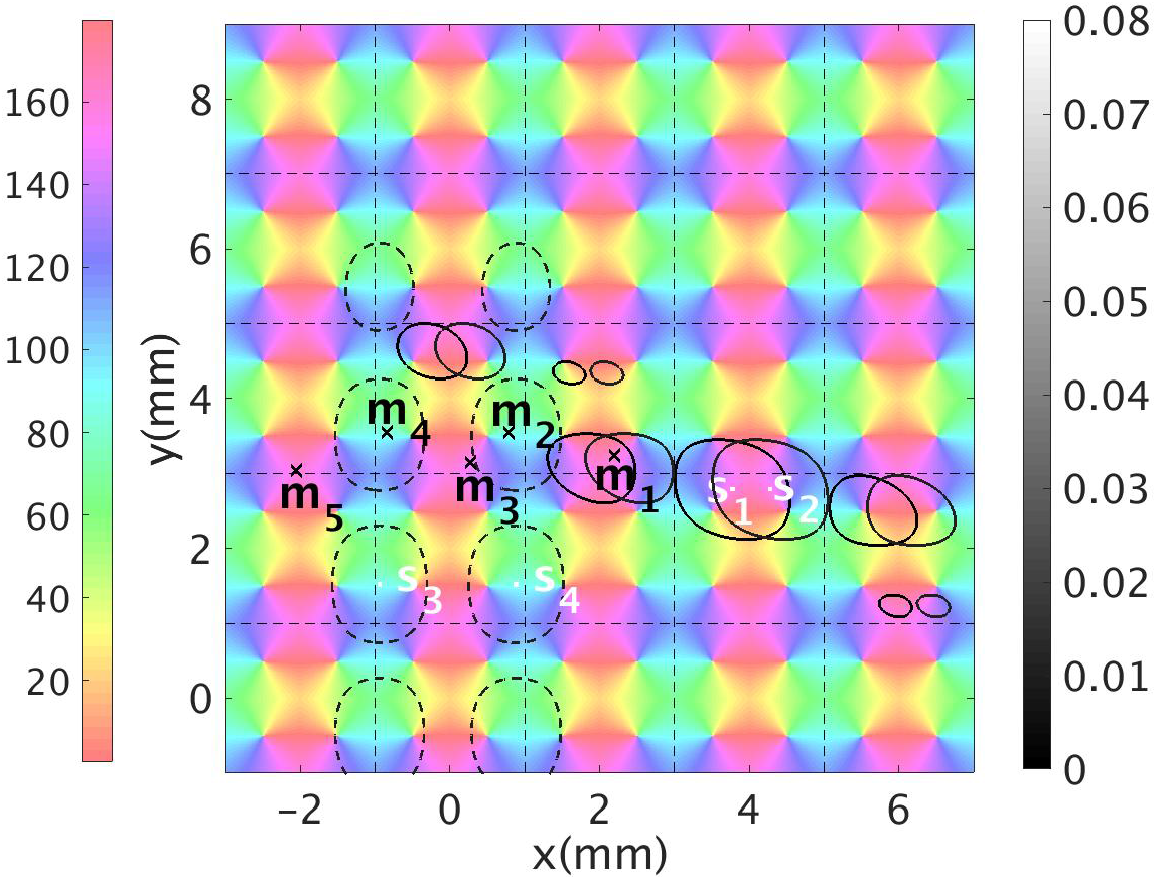
Schematic of the two experimental conditions, showing measurement points **m**_1_ to **m**_5_ and source points on V1. The first experimental condition corresponds to a stimulus at 157° with corresponding sources denoted **s**_1_ and **s**_2_, with solid gray contours showing the propagation. The second experimental condition corresponds to the a 90° stimulus. Here, **s**_3_ and **s**_4_ are the sources and dotted gray contours show the propagation strength according to the grayscale at right. Dotted vertical and horizontal lines bound unit cells and the color bar shows OP in degrees.

### 7.3 Comparison of Predicted and Experimental Correlation Functions

According to the experimental findings in Engel et al. (1990), when the input light bar is oriented at 157.5°, measurement sites **m**_1_, **m**_3_, and **m**_5_ have synchronized oscillatory responses; and, when the input light bar is oriented at 90°, **m**_2_ and **m**_4_ are stimulated simultaneously. Figure 10 shows the CCFs and ACFs calculated from the experimental data. In Figure 10(a), the synchronized activities at **m**_1_, **m**_3_, and **m**_5_ are evoked by a 157.5° oriented stimulus. All the cross correlograms are peaked at zero time-lag and have an average oscillation frequency of ~54 Hz. The envelope of the correlograms decreases to 1/*e* of its center peak value at around 45 ms. The ACFs and CCF of **m**_2_ and **m**_4_ from a vertical light bar stimulus are shown in Figure 10(b). The CCF between **m**_2_ and **m**_4_ oscillates at around 55 Hz, and it takes more than 50 ms for the correlation strength to decrease to 1/*e* of its maximum.

**Figure 10:**
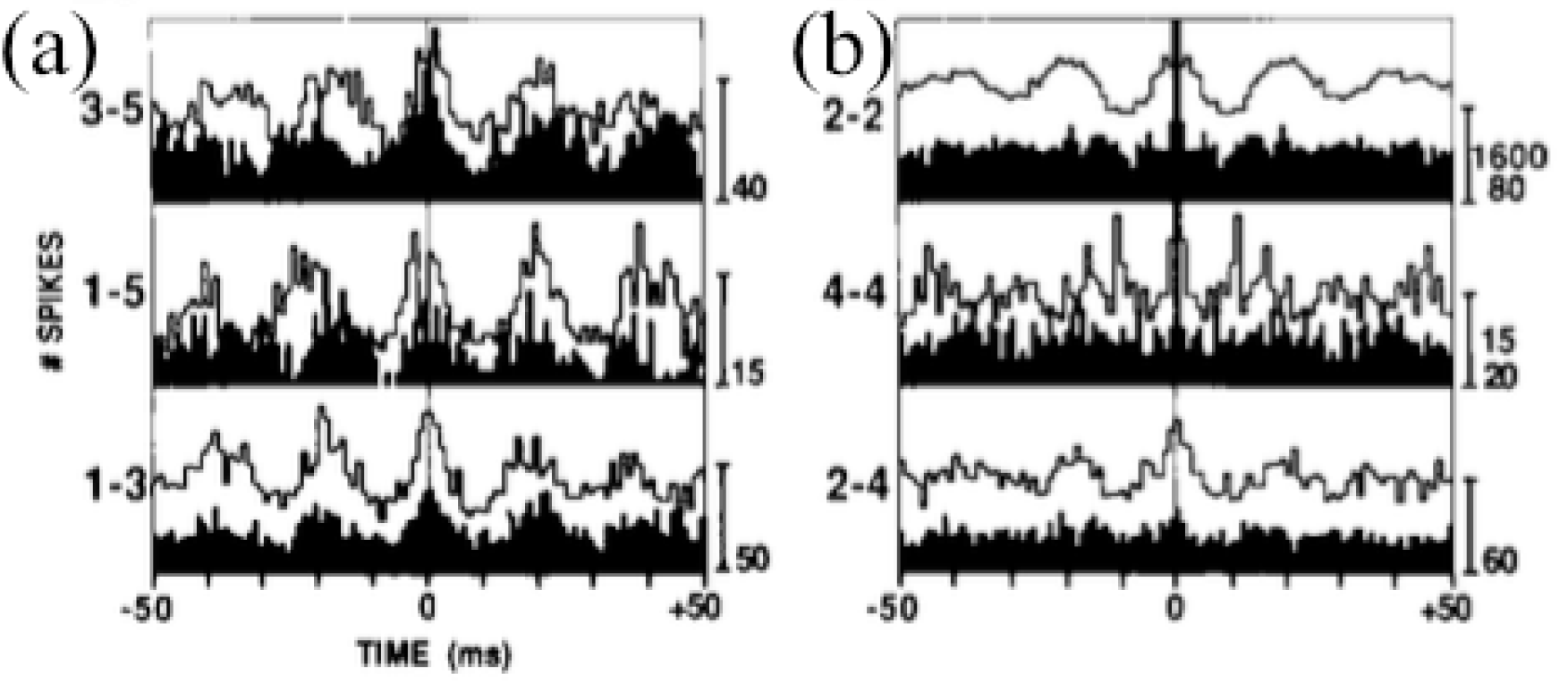
Cross correlograms from experiments recordings calculated by Engel et al. (1990). In each case a baseline level of activity shifts the oscillatory part of the correlation upward and must be subtracted for comparison with the theoretical results. (a) Cross correlograms between measurement sites **m**_3_ and **m**_5_, **m**_1_ and **m**_5_, and **m**_1_ and **m**_3_ corresponding to an input light bar oriented at 157.5°. (b) Auto correlograms of **m**_2_ and **m**_4_ in the top two row, and cross correlograms on the bottom row between **m**_2_ and **m**_4_, corresponding to the vertical light bar.

Moreover, in the experiments it was also found that the correlation strength between **m**_1_ — **m**_5_ is weaker than that between **m**_1_ — **m**_3_ and between **m**_3_ — **m**_5_ (i.e., the bar that indicates the number of spikes, on the right of the plot in the second row of Figure 10(a), has smaller number than other two plots). This is due to the fact that the spatial distance between **m**_1_ and **m**_5_ is the largest, and the correlation strength falls off with distance.

We next explore the properties of our predicted correlation functions using Eq. (55) with the experimental conditions. Figure 11(a) shows the plots of our predicted temporal correlation functions between **m**_3_ and **m**_5_, **m**_1_ and **m**_5_, and **m**_1_ and **m**_3_. Similarly to the experimental CCFs, all the theoretical CCFs: (i) are oscillatory and peak at zero time lag; (ii) have an oscillation frequency around 57 Hz; and, (iii) have their characteristic time for the correlation envelope to decrease by 1/*e* of the maximum value at approximately 40 ms. These theoretical results agree with the experimental results, once a nonzero mean baseline is subtracted from the latter.

Our prediction also captures the spatial dependence of the maximum correlation strength. The plot in the middle row of Figure 11(a) corresponds to the correlation between **m**_1_ — **m**_5_ and has the smallest amplitude among the three CCFs.

**Figure 11:**
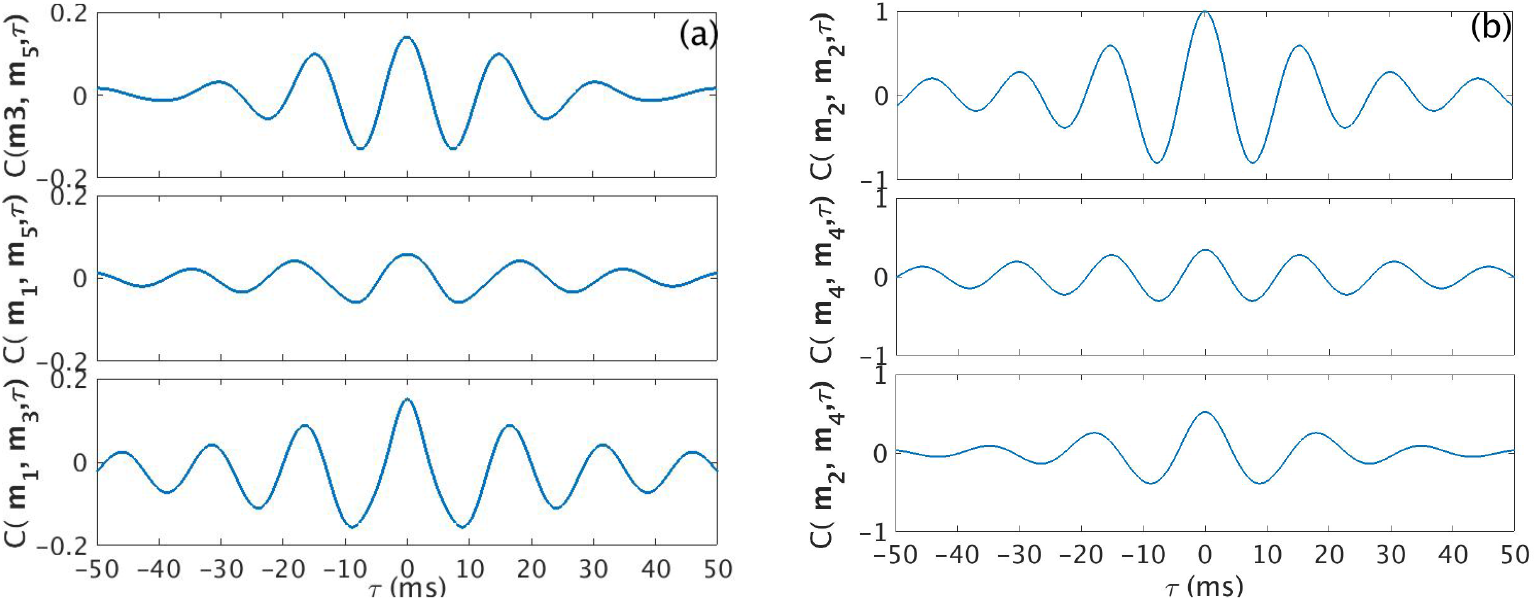
Normalized temporal cross correlation with zero mean for experimental conditions with stimuli at 157.5° and 90°. (a) Normalized temporal correlation between measurement sites **m**_3_ and **m**_5_, **m**_1_ and **m**_5_, and **m**_1_ and **m**_3_ for a stimulus at 157.5°. Such condition is illustrated in Figure 9. (b) Normalized temporal cross correlation between **m**_2_ and **m**_4_, and the autocorrelation function at **m**_2_ and **m**_4_, for a stimulus at 90°.

Figure 11(b) shows the predicted temporal correlation function generated by the vertical input light bar. In order to be consistent with the experimental results shown in Figure 10(b), the autocorrelation functions of **m**_2_ and **m**_4_ are also included in the top two rows of Figure 11(b). Both ACFs show oscillations in the gamma band. The CCF between **m**_2_ and **m**_4_ shows a center peak at *τ* = 0 and oscillates at 55 Hz. The time for the envelope decay to 1/*e* of the center peak value is 25 ms. These properties are also in line with the experimental findings.

## 8 Summary and Conclusion

We have generalized the spatiotemporal correlation functions in two dimensions that incorporates the spatial structure of the OP map and OD columns of V1. Our results show that the neural activities are synchronized in gamma band when neurons have similar feature preference. The main results are:

i. The derivation of a shape function that modulates the spatial patchy propagation of the neural signals. The shape function models the propagation such that the orientation of the propagation direction is aligned with the OP of source, and the connected neurons are patchy and periodically located. The parameters of the shape function are tuned to match the propagation ranges observed in experiments (Bosking et al., 1997).
ii. The systematic characterization of the 2D two-point temporal correlation function. The generalized correlation function is evaluated numerically for various combinations of stimulation and measurement sites. The results demonstrate a synchronized gamma oscillation exists between two groups of neurons that have similar OP to the sources. The correlation strength is larger for inter-columnar connections than for intra-columnar connections. As the measurement points are further away from the sources, the correlation strength decreases, and is negligible when the spatial separation of the measurement points exceeds 7 mm.
iii. The construction of a 2D correlation maps. These maps show the changes expected in the peak correlation strength with respect to the variation of the OP of one of the measurement sites, and its distance to a second measurement site. The positive correlations appear as patches on an axis oriented at the OP of the source; and, negative correlations occur where the OPs of the measurement sites are orthogonal to the OP of the source.
iv. The comparison of the predicted temporal correlations using experimental conditions. Our theoretical results are compared with the experimental findings and shows there is a close match between both in terms of the oscillation frequency and the characteristic decay time of the correlation function envelope. In addition, our CCFs also capture the spatial dependence of correlation strength, which decreases with distance between the measurement sites.

Overall, our generalized spatiotemporal correlation function reproduces the gamma band oscillations observed in V1 and relates the spatially distributed neural responses to the periodic spatial structure of OP and OD in V1. This study lays the foundation to further investigate other visual perception phenomena such as the binding problem.

Future work will focus on using a more realistic lattice of pinwheels and introduce asymmetries between the left/right OD columns to account for strabismus.

## 9 Acknowledgements

This work was supported by an Australian Research Council Laureate Fellowship (grant FL1401000025) and the Australian Research Council Center of Excellence for Integrative Brain Function (grant CE140100007).

